# A transformation of oxygen saturation (the saturation virtual shunt) to improve clinical prediction model calibration and interpretation

**DOI:** 10.1101/391292

**Authors:** Guohai Zhou, Walter Karlen, Rollin Brant, Matthew Wiens, Niranjan Kissoon, J Mark Ansermino

**Affiliations:** Department of Statistics, University of British Columbia, Vancouver, Canada; Department of Health Sciences and Technology, ETH Zurich, Zurich, Switzerland; School of Population and Public Health, University of British Columbia, Vancouver, British Columbia, Canada; Department of Pediatrics, University of British Columbia, Vancouver, Canada; Department of Anesthesiology, Pharmacology & Therapeutics, University of British Columbia, Vancouver, Canada

**Keywords:** Pediatrics, Public Health, Improvement Science, Oxygen Saturation, Shunt, Transformation

## Abstract

**Background:** The relationship between peripheral oxygen saturation (SpO_2_) and the inspired oxygen concentration is non-linear. SpO_2_ is frequently used as a dichotomized predictor, to manage this non-linearity. We propose the *saturation virtual shunt (VS)* as a transformation of SpO_2_ to a continuous linear variable to improve interpretation of disease severity within clinical prediction models.

**Method:** We calculate the *saturation VS* based on an empirically derived approximation formula between physiological VS and SpO_2_. We evaluated the utility of the *saturation VS* in a clinical study predicting the need for facility admission in children in a low resource health-care setting.

**Results:** The transformation was *saturation VS = 68.864*log_10_(103.711* − *SpO_2_) −52.110*. The ability to predict hospital admission based on a dichotomized SpO_2_ produced an area under the receiver operating characteristic curve of 0.57, compared to 0.71 based on the untransformed SpO_2_ and *saturation VS*. However, the untransformed SpO_2_ demonstrated a lack of fit compared to the *saturation VS* (goodness-of-fit test p-value <0.0001 versus 0.098). The observed admission rates varied non-linearly with the untransformed SpO_2_ but varied linearly with the *saturation VS.*

**Conclusion:** The *saturation VS* estimates a continuous linearly interpretable disease severity based on SpO_2_ and improves clinical prediction.

## INTRODUCTION

Parenchymal lung disease results in decreased efficiency of gas exchange in the lung and manifests as hypoxaemia [1]. For clinicians, the peripheral oxygen saturation (SpO_2_) is widely used as a measure of the degree of lung impairment. However, the well-known non-linear hemoglobin-oxygen dissociation curves prohibit the utility of SpO_2_ as a *linear* marker of disease severity and clinical outcomes such as the need for hospital admissions.

To circumvent this limitation, we propose a simple transformation of SpO_2_, the *saturation virtual shunt* (VS), which is *linearly* related to the degree of impairment in oxygen exchange and that can be adjusted for the fraction of inspired oxygen (FiO_2_). This transformation is based on the known empiric and physiological relationship between VS, a theoretical concept that describes the overall loss of oxygen content between the inspired gas and the arterial blood [2], and SpO_2_. This transformation is aimed at addressing the non-linearity in hemoglobin-oxygen dissociation curves for use in prediction models.

The concept of VS was initially used to describe the non-linear relationship between FiO_2_ and the arterial partial pressure of O_2_ (PaO_2_) using iso-shunt curves [2]. VS can be defined as the proportion of blood that would need to bypass the lungs to produce the difference between the calculated end capillary venous oxygen content and arterial blood oxygen content and is also known as venous admixture [2]. Estimating VS is useful as the magnitude of VS quantitatively describes the efficiency of the gas exchange and the severity of a disease process that may lead to hypoxaemia.

VS is typically calculated based on assumed values of the arterial/mixed venous oxygen content difference, hemoglobin level, pH and temperature unless these values can be measured [3, 4]. A more common formula, the difference between the oxygen partial pressure in the alveoli (PAO_2_) and systemic arteries (PaO_2_) (P[A-a] O_2_), has been used to represent the shunt calculation. This formula does adjust for changes in FiO_2_ (due to changes in altitude or oxygen administration) but the reliance on an invasive measurement to obtain PaO_2_ makes it impractical and the partial pressure of oxygen is used as a surrogate for oxygen content. The partial pressure of oxygen however is not linearly related to the degree of gas exchange abnormality and hence this approach is sub-optimal [3–5].

The objectives of this study are the following: (1) to use the known physiological relationship between VS and SpO_2_ to derive the *saturation VS* as a transformation of SpO_2_ (for clinical interpretation and prognostic research) and (2) to evaluate the *saturation VS* as a predictor of hospital admission, compared to the dichotomized SpO_2_ and the untransformed SpO_2_, in a cohort of children visiting the emergency department at the Kamudini Women Medical College Hospital in Bangladesh [6].

## METHODS

### Physiological calculation of VS

Following Karlen W, et al [3, 4], we derive physiological VS from inspired oxygen FiO_2_ and arterial oxygen saturation (SaO_2_) for *a full range of theoretical subjects* that satisfy the following assumptions.

- The loss due to capillary diffusion is negligible, which allows alveoli oxygen content (PAO_2_) to be approximated by end-capillary oxygen content.
- SaO_2_ is estimated without error from SpO_2_ obtained from pulse oximetry.
- Patients are on room air at the time of SpO_2_ measurements. The barometric pressure is at sea level (101 kPa) and inspired oxygen (FiO_2_) is at 21%.
- Values for water vapor pressure, alveolar CO_2_ partial pressure, respiratory quotient, incomplete capillary diffusion, arterio-venous oxygen difference, oxygen-binding capacity of hemoglobin, blood concentration of hemoglobin, and solubility of O_2_ in hemoglobin are assumed to be normal and constant.

With the above assumptions, we can *theoretically* calculate the physiological VS using the previously established Alveolar Gas Equation and the Severinghaus and Severinghaus-Ellis equations [7–9]. PAO_2_ was estimated using the Alveolar Gas Equation [7]. Alveoli oxygen concentration (SAO_2_) was then estimated from PAO_2_ using the Severinghaus equation [8]. To calculate arterial oxygen content, SaO_2_ was transformed to PaO_2_ using the Severinghaus-Ellis equation [8, 9]. The detailed mathematical descriptions of the above calculation can be found in the online supplementary material or in Karlen W, *et al* [3, 4].

### Saturation virtual shunt

To produce a simple and more clinically useful method to describe the non-linear relationship between the physiological VS and SpO_2_, we fitted several common non-linear functions, such as polynomials and logarithmic functions. The unknown parameters of these functions were estimated using the non-linear least squares method [10]. For this fitting process, we selected SpO_2_ at 1% intervals in the range from 50% to 98%. We chose this range of values because the previously described empiric formulae are not valid for SpO_2_ values above 98% [3, 4]. We also excluded SpO_2_ values below 50% as they are rare and typically associated with severe clinical cyanosis.

The *saturation VS* was then defined based on the fitted relationship between the physiological VS and SpO_2._

### Application to Prediction Performance

We evaluated the use of the *saturation VS* compared to the dichotomized SpO_2_ and the untransformed SpO_2_ in a recently completed prospective observational study at the Kumudini Women’s Medical College Hospital’s in Bangladesh, a rural tertiary care hospital [6]. Ethics approval and informed consent were obtained prior to data collection.

The study aimed to develop a simple model to predict the need for facility admission that could be used in a community setting. Children < 5 years of age presenting at the outpatient or emergency department were enrolled. Study physicians collected clinical signs and symptoms from the facility records and performed recordings of SpO_2_, heart rate and respiratory rate. Facility physicians made the decisions about the need for hospital admission on clinical grounds without knowledge of the oxygen saturation measurements. SpO_2_ value was taken as the median over a minute at the time of initial assessment. SpO_2_ readings above 98% were considered to be equal to 98%, because 98% is the theoretical maximum reading possible on room air at sea level. Readings above 98% occurred due to the tolerance level or bias of the pulse oximeters [3, 4]. Children who showed high SpO_2_ variability (range>6%) in combination with low perfusion were excluded. Motion artifact, ambient light and poor positioning of the sensor typically resulted in high variability and low perfusion leading to a high likelihood of erroneous SpO_2_ readings. Low perfusion was assessed post-hoc by based on the amplitude of the photoplethysmogram and the pulse oximeter device perfusion index (low/medium/high). Children with SpO_2_ below 75% (a danger sign) were also excluded from predictive modeling, for they were considered critically ill and should be directly admitted into higher-level facilities regardless of any model-based predictions [11].

We fitted three univariate logistic regression models with different predictors to predict the need for facility admission: 1) hypoxaemia, defined as SpO_2_ <90% by the World Health Organization (WHO, dichotomized SpO_2_ model), 2) the observed SpO_2_ (untransformed SpO_2_ model), and 3) the *saturation VS* (saturation VS model). We compared these models in terms of overall accuracy, calibration and clinical interpretation. Overall accuracy was assessed using the area under the receiver operating characteristic curve (AUC ROC) and its 95% confidence interval (CI). AUC ROC measured the probability that a randomly selected admitted child would receive a higher predicted probability of requiring admission than a randomly selected child who was not admitted [12]. For the untransformed SpO_2_ model and the saturation VS model, calibration was assessed by plotting the observed admission rate against the group average of the predicted probability of requiring admission for each of 3 groups determined a priori: SpO_2_ <90%, SpO_2_ from 90% to 97.5% and SpO_2_ equal to 98%. A chi-square goodness-of-fit test was then applied [13]. In addition, the observed admission rates were plotted against 10 equally spaced SpO_2_ or 10 equally spaced *saturation VS* categories sharing similar ranges. The plots were fitted to linear and non-linear trends using the method of least squares [10], which aimed to minimize the total squared difference between the observed admission rates and the risks of admission directly interpreted from the category-average SpO_2_ or saturation VS levels. The accuracy of the fitted relationship was quantified by the standard deviations of the difference between the observed admission rates and the interpreted risks of admission based on the SpO_2_ or *saturation VS* category labels. The 95% CIs for these standard deviations were calculated based on chi-square distributions [10].

## RESULTS

### Physiological Calculation of Virtual Shunt

Due to the empirical nature of the physiological equations [7–9], the VS corresponding to SpO_2_ 98% was a small negative value (−0.78). We thus added 0.78 to all VS values so that a normal SpO_2_ 98% corresponded to exactly zero VS.

### The Transformation Formula for the Saturation Virtual Shunt

The functions of the form y = a*log_10_(b − x) + c were sufficient to capture the non-linearity in the relationship between physiological VS and SpO_2_. The relationship was statistically best fitted by an equation *VS* = *68.864*log*_*10*_*(103.711* − *SpO*_*2*_) −*52.110* (Figure 1). The *saturation VS* was therefore defined as *saturation VS* = *68.864*log_10_(103.711* − *SpO_2_) −52.110*.

**Figure 1:**
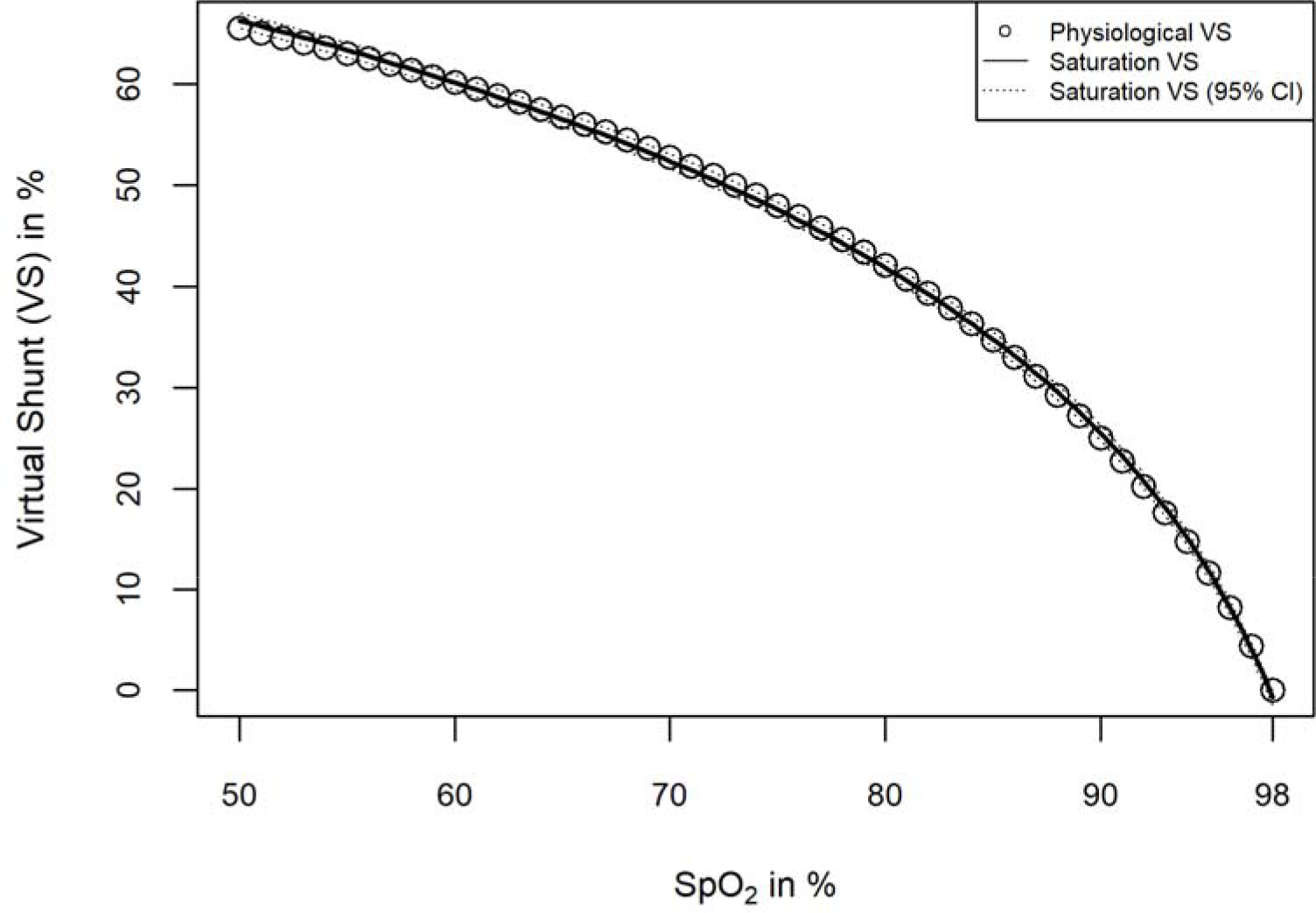
Scatterplot of the physiological virtual shunt (%) and the *saturation VS* (%) against SpO_2_ (%). “Physiological VS” is the physiological virtual shunt estimated by solving simultaneous equations using multiple physiological variables, “Saturation VS” is computed by “Saturation VS = *68.864*log_10_(103.711* − *SpO_2_) −52.110*”. The standard deviation of the differences between “Physiological VS” and “Saturation VS” is 0.37%, indicating that 95% of the differences between “Saturation VS” and “Physiological VS” are within 0.74%.

### Prediction Performance and Model Calibration and Interpretation

In total, 2943 of the 3374 recruited cases had adequate SpO_2_ recordings, of whom 831 were admitted and 2112 were not admitted. We excluded 5 cases showing SpO_2_ variability > 6% in combination with low perfusion. We adjusted the 868 SpO_2_ readings above 98% to be equal to 98%. The 12 cases with SpO_2_ <75% were all admitted and they were excluded from predictive modeling.

The distribution of the untransformed SpO_2_ and that of the *saturation VS* revealed more informative discrimination of the outcome group than that of the dichotomized SpO_2_ (Figure 2). In addition, the distribution of the *saturation VS* was less skewed (skewness 2.26 vs −3.54 for the admitted and 0.98 vs −2.84 for the not admitted cases) than that of the untransformed SpO_2_ (Figure 2).

**Figure 2:**
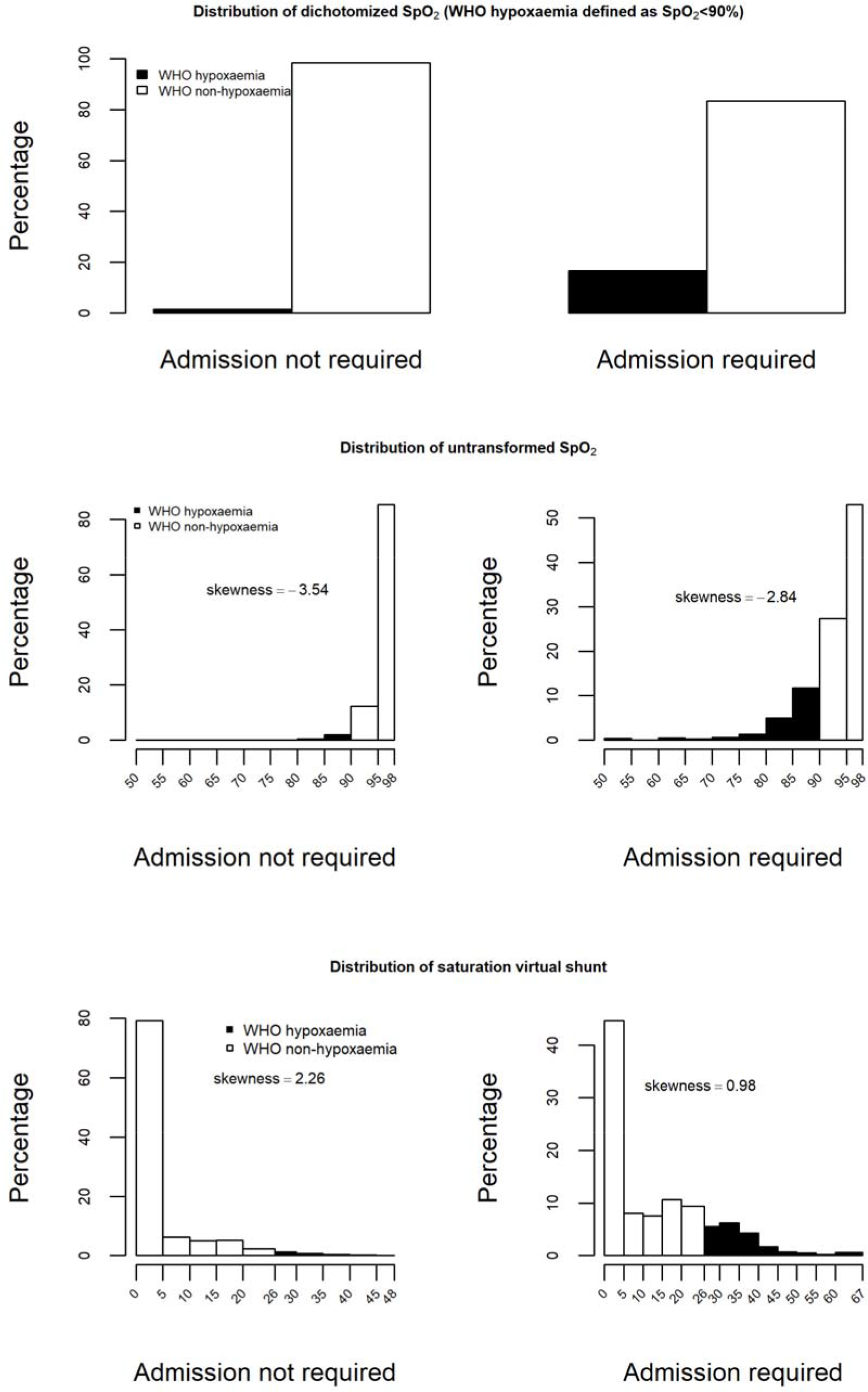
Distribution of dichotomized SpO_2_, untransformed SpO_2_ (%) and the *saturation VS* (%) by outcome group. By having a less skewed distribution, the *saturation VS* is superior to the untransformed SpO_2_ in terms of the ability to more evenly stratify patients by sickness severity.

The dichotomized SpO_2_, the untransformed SpO_2_ model and the *saturation VS* model all demonstrated that a SpO_2_ below 90% was associated with increased risk of admission and the latter two, unsurprisingly had a much higher AUC ROC (Table 1). Despite the identical AUC ROC, the untransformed SpO_2_ model demonstrated a statistically significant lack of fit (p-value <0.0001, χ^2^ =19.973, df=1) whereas the saturation VS model did not (p-value =0.098, χ^2^ =2.744, df=1). A closer look at the data revealed that the untransformed SpO_2_ model significantly underestimated the risk of admission among the 1017 children with SpO_2_ between 90% and 97.5% and significantly overestimated the risk of admission among the 1750 children with SpO_2_ >=98% (Figure 3). Therefore, the saturation VS model was better calibrated than the untransformed SpO_2_ model. In terms of clinical interpretation, a 5% decrease in SpO_2_ and a 5% increase in the saturation VS were respectively predicted to be associated with a 286% and 55% increase in the odds of requiring admission (Table1). The magnified odds ratio obtained from the untransformed SpO_2_ model was due to the dense distribution of SpO_2_ data between 85% and 98% (Figure 2) and that in this region a 5% decrease in SpO_2_ corresponded to a more than 5% increase in the *saturation VS* (Figure 1).

**Table 1:** Summary of the three prediction models for the need of facility admission

| **Model** | **Odds ratio (95% CI)** | **AUC ROC (95% CI)** |
| --- | --- | --- |
| Dichotomized SpO<sub>2</sub> model | 11.47 (7.74-16.99)<sup>a</sup> | 0.57 (0.56-0.58) |
| Untransformed SpO<sub>2</sub> model | 3.86 (3.31-4.50)<sup>b</sup> | 0.71 (0.69-0.73) |
| Saturation VS model | 1.55 (1.48-1.62)<sup>c</sup> | 0.71 (0.69-0.73) |

a. associated with changing from the absence to presence of WHO defined hypoxaemia
b. associated with a 5% decrease in SpO<sub>2</sub>
c. associated with a 5% increase in the *saturation VS*

**Figure 3:**
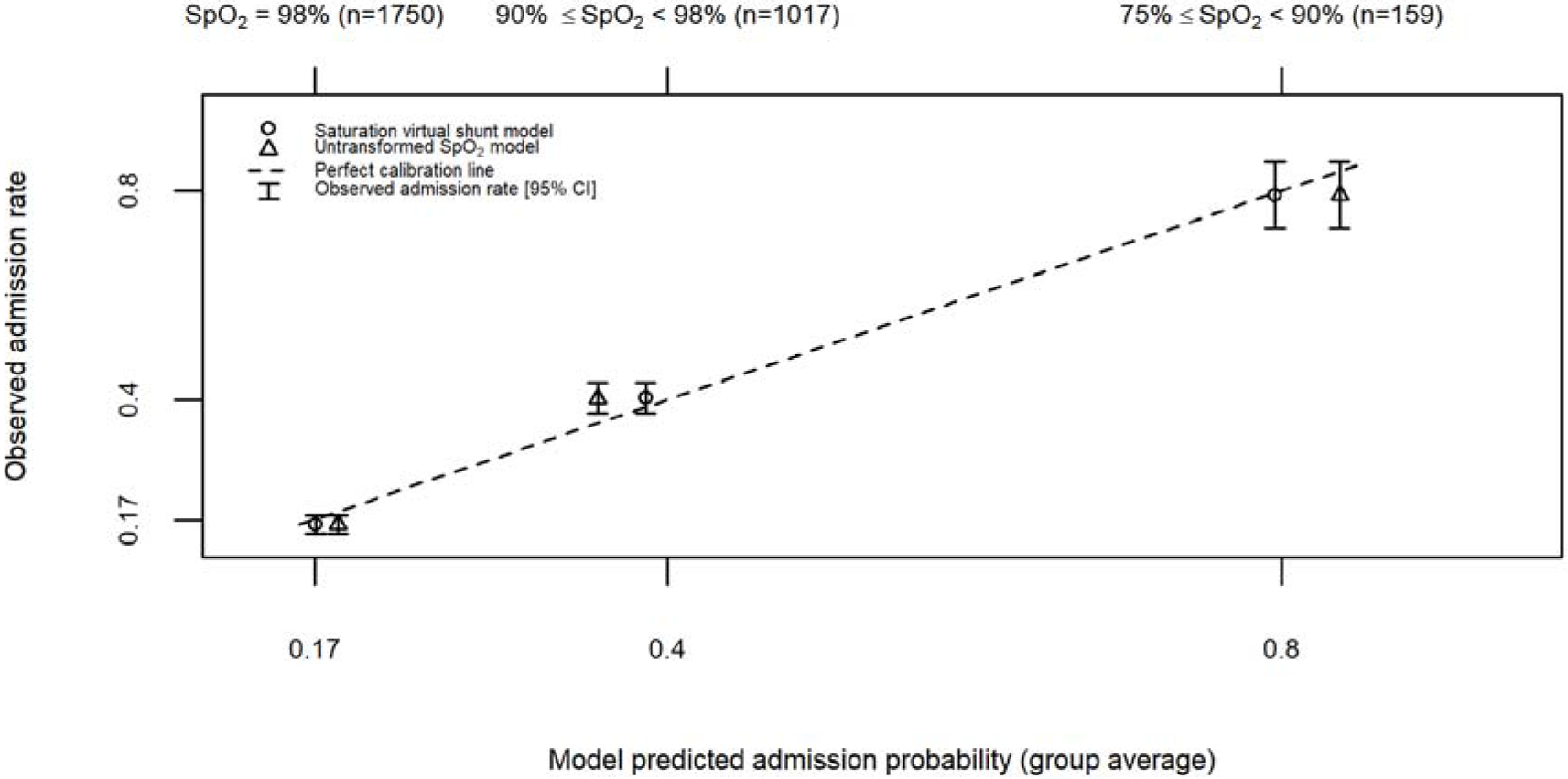
Calibration plot of the untransformed SpO_2_ model and the saturation VS model applied to the 2926 cases with SpO_2_ above or equal to 75%. The dotted line is the line of equality on which model predicted admission probabilities perfectly coincide with observed admission rates.

The observed admission rates exhibited a non-linear relationship with SpO_2_ but an approximately linear relationship with *saturation VS* (Figure 4). More specifically, each 4% increase in the *saturation VS* was on average associated with an approximately 8.2% increase in the admission rate (e.g., 286 out of 1757 children were admitted with the *saturation VS* from 0% to 4%, whereas 84 out of 288 children were admitted with the *saturation VS* from 12% to 16%). In contrast, each 2% decrease in SpO_2_ would be associated with *varying* increases in the admission rate due to the nature of the non-linear trend in Figure 4.

**Figure 4:**
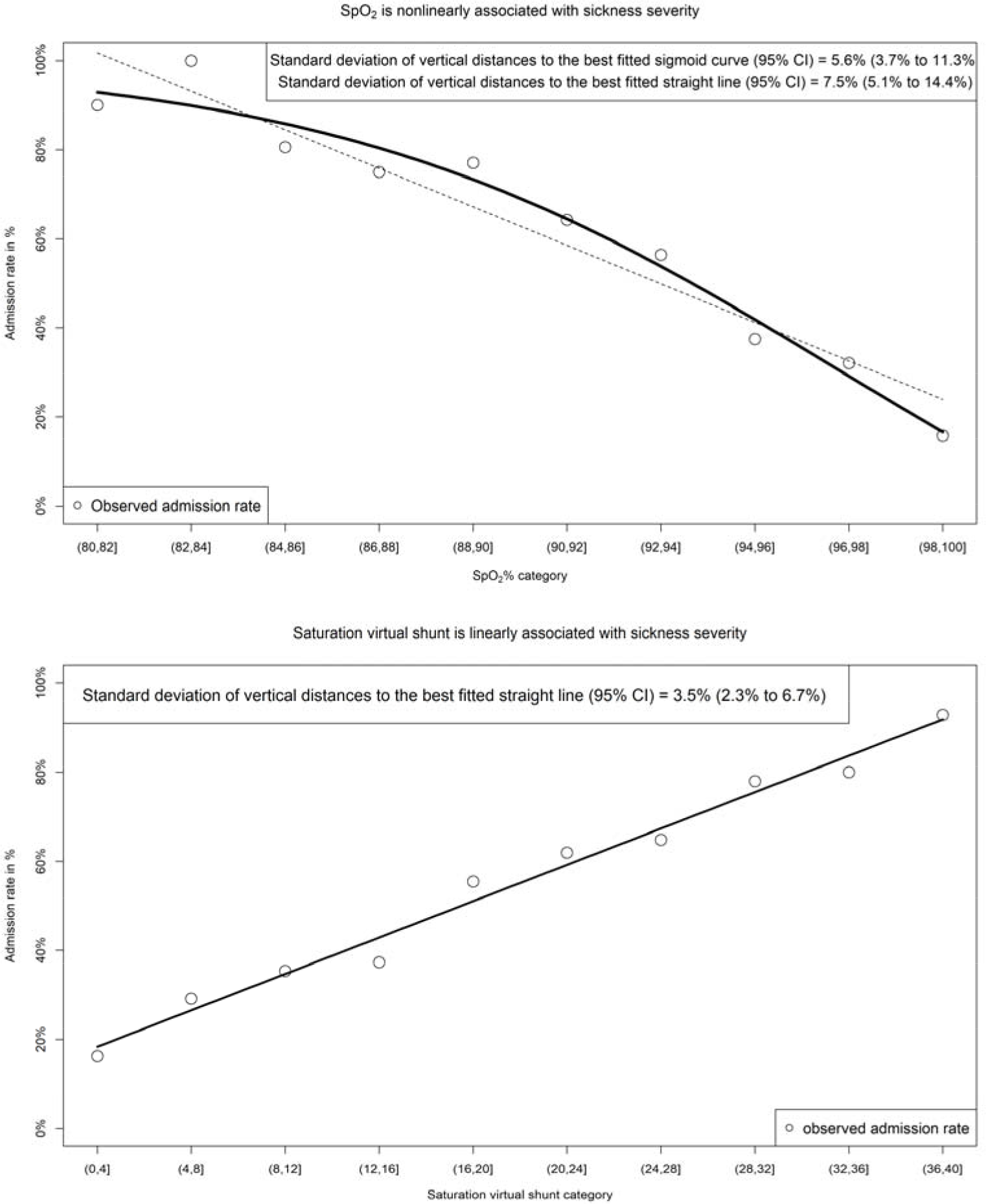
Observed admission rates compared to equally spaced SpO_2_ and *saturation VS* categories. Each category includes the right endpoint but excludes left endpoint (e.g. 80-82 includes SpO_2_ =82% but excluded SpO_2_ =80%). For the curve-fitting, the categories are coded from 1 to 10. For SpO_2_, the best fitted sigmoid curve (a common type of non-linear curves for S-shaped relationships) corresponds to an equation y = 0.980 − 2.138/(1+ e^−0.266*(x−11.073)^).

## DISCUSSION

The transformation of the SpO_2_ to the *saturation VS* improves clinical interpretation, accuracy and calibration of prediction models. For instance, in our study in under 5 children in Bangladesh the *saturation VS* improved clinical prediction, calibration and interpretation of the need for hospital admissions. This is not surprising because dichotomizing the SpO_2_ is ill-suited to decision making in clinical medicine and resulted in a significant degradation of prediction performance. The *saturation VS* may also have additional importance in estimating severity of disease and response to treatment (such as oxygen administration) since it incorporates the non-linearity in hemoglobin-oxygen dissociation curves. Small changes in SpO_2_ on the flat portion of the oxygen saturation curve (near 100%) reflect a much greater change in physiology than the same change at a lower SpO_2_. In contrast, the saturation VS is *linearly* related to the changes in the physiological and clinical state and provides more granular information of adverse changes in physiology. For example, on the scale of SpO_2_, a decrease from 95% to 90% would reflect more impairment in gas exchange, and therefore may indicate more significant change (deterioration in clinical condition), than a decrease from 90% to 85%. This non-linear interpretability of SpO_2_ is particularly undesirable when it is used as a predictor (e.g. in Amatet, *et al* [14]), whether in univariate or multivariate analysis, for interpretations of regression models often involve a description of the average amount of outcome change that will be associated with a given amount of predictor change. Such description is only meaningful if the amount of predictor change is *clinically comparable* for different baseline values. This has typically been resolved by dichotomizing the SpO_2_ values. However, the use of continuous predictors has been recommended to prevent information loss and decrease in predictive capability resulting from dichotomization [15]. To improve model interpretability, it would be unwise to use hypoxaemia as a surrogate predictor in view of the loss in accuracy. Instead, the use of the *saturation VS* in lieu of observed SpO_2_ as a predictor would not only maintain the prediction accuracy but also increase clinical interpretation and calibration of prediction models. Such benefits are especially valuable for resources-limited settings where staff trainings may also be inadequate. The use of equally spaced saturation VS categories may also provide a more intuitive interpretation for clinicians to *linearly* interpret sickness severity (e.g. hospital admission rate) that is not achievable with the direct use of the untransformed SpO_2_.

The major limitation of the proposed transformation is that the derivation makes many assumptions about normal clinical conditions. A change in the *saturation VS* may be a result of changes in other unmeasured variables in the model and may not be a result of abnormal gas exchange. A further limitation is that this study modeled the outcome of admission, which was not necessarily linked to issues of respiratory compromise. This would be artificially associated with lower AUC values than if modeled using a cohort of children being assessed with a presumed respiratory illness. However, since our predictive variables were all based on oxygen saturation, the comparative differences remain internally valid. Despite these limitations, this approach has the potential to fill an important gap in the utilization of oxygen saturation data in both clinical and research settings. Further validation in clinical settings is therefore required to better define it utility in this context.

In conclusion, the SpO_2_ transformed *saturation VS* provides an intuitive measure of hypoxaemia and may prove to be a useful aid in clinical practice when measuring SpO_2_ and as a component of clinical prediction models when included with electronic devices (such as mobile phones) that can easily perform the required calculation. Further validation is necessary prior to adoption into clinical practice.

## ACKNOWLEDGMENTS

We would like to acknowledge Charles Larson (The University of British Columbia) and Shams El Arifeen (Centre for Child and Adolescent Health, International Centre for Diarrhoeal Disease Research, Bangladesh, Dhaka, Bangladesh) and their teams involved in the Interrupting Pathways to Sepsis Initiative for the permission to use the clinical data.

## AUTHOR CONTRIBUTIONS

JMA, RB and NK contributed to the design of the study. GZ, WK and RB analyzed the data. GZ, WK, RB, MW, JMA and NK wrote, revised and approved the manuscript.

## STATEMENT OF FINANCIAL SUPPORT

This study was a component of the project titled Interrupting Pathways to Maternal, Newborn and Early Childhood Sepsis (IPSI), funded by the MUSKOKA Initiative on Maternal, Newborn and Child Health (MNCH).

## DISCLOSURE STATEMENT

The authors confirm that there are no competing interests.

